# Robust and sensitive detection of SARS-CoV-2 using PCR based methods

**DOI:** 10.1101/2020.07.03.186304

**Authors:** Changwoo Park, Jina Lee, Zohaib ul Hassan, Keun Bon Ku, Seong Jun Kim, Hong Gi Kim, Edmond Changkyun Park, Gun-Soo Park, Daeui Park, Seung-Hwa Baek, Dongju Park, Jihye Lee, Sangeun Jeon, Seungtaek Kim, Chang-Seop Lee, Hee Min Yoo, Seil Kim

## Abstract

The World Health Organization (WHO) has declared the Coronavirus disease 2019 (COVID-19) as an international health emergency. Current diagnostic tests are based on the reverse transcription-quantitative polymerase chain reaction (RT-qPCR) method, the gold standard test that involves the amplification of viral RNA. However, the RT-qPCR assay has limitations in terms of sensitivity and quantification. In this study, we tested both qPCR and droplet digital PCR (ddPCR) to detect low amounts of viral RNA. The cycle threshold (CT) of viral RNA by RT-PCR significantly varied according to the sequence of primer and probe sets with in vitro transcript (IVT) RNA or viral RNA as templates, whereas the copy number of viral RNA by ddPCR was effectively quantified with IVT RNA, cultured viral RNA, and RNA from clinical samples. Furthermore, the clinical samples were assayed via both methods, and the sensitivity of the ddPCR was determined to be significantly higher than RT-qPCR. These findings suggest that ddPCR could be used as a highly sensitive and compatible diagnostic method for viral RNA detection.

## Introduction

Coronaviruses have caused several notable respiratory disease outbreaks in humans, such as severe acute respiratory syndrome (SARS). The SARS coronavirus was the first fatal pathogenic human coronavirus to be identified and caused 8,096 infections and 774 deaths in 26 countries (“WHO | Summary of probable SARS cases with onset of illness from 1 November 2002 to 31 July 2003,” 2015). Prior to the emergence of SARS-CoV, only two HCoVs (HCoV-229E and HCoV-OC43) were known, both of which caused mild respiratory symptoms (Fung & Liu, 2019). Bats were the natural hosts of SARS-CoV and civet cats were regarded as the intermediate host (Li et al., 2006). Extensive studies on bat coronaviruses showed that such coronaviruses could be potential human pathogens (Ge et al., 2013; Wu, Peng, Wilken, Geraghty, & Li, 2012). MERS-CoV was first reported in 2012 and caused 2,494 infections and 858 deaths in 27 countries as of November 2019 (“WHO | Middle East respiratory syndrome coronavirus (MERS-CoV),” in press). MERS-CoV also originated from bat coronaviruses and camels were the intermediate host (Ashour, Elkhatib, Rahman, & Elshabrawy, 2020).

According to the World Health Organization (WHO), the WHO China Country Office was informed of cases of pneumonia of unknown etiology in Wuhan City, Hubei Province, on December 31st, 2019 (World Health Organization (WHO), 2020). The novel coronavirus, currently termed SARS-CoV-2, was officially announced as the causative agent by the International Committee on Taxonomy of Viruses (ICTV) (Coronaviridae Study Group of the International Committee on Taxonomy of Viruses, 2020; Gorbalenya, 2020). SARS-CoV-2, or severe acute respiratory syndrome coronavirus 2, has resulted in 2,114,269 laboratory-confirmed cases and 145,144 deaths as of April 17th, 2020 (da Costa, Moreli, & Saivish, 2020). A viral genome sequence was released for immediate public health support via the community online resource virological.org on January 10th (Wuhan-Hu-1, GenBank accession number MN908947) (Li et al., 2006), followed by four other genomes that were deposited on January 12th in the viral sequence database curated by the Global Initiative on Sharing All Influenza Data (GISAID). The genome sequences suggest the virus is closely related to the members of a viral species termed severe acute respiratory syndrome (SARS)-related CoV, a species defined by the agent of the 2002/03 outbreak of SARS in humans (Ge et al., 2013; Wu et al., 2012). The species also comprises of a large number of viruses mostly detected in rhinolophid bats in Asia and Europe.

For the past several decades, quantitative polymerase chain reactions (qPCR) have become the gold standard for quantifying relative gene expressions. On the other hand, the recently developed digital polymerase chain reaction (dPCR) enables the highly sensitive measurement and absolute quantitation of nucleic acids (Hindson et al., 2011, 2013). dPCR does not require calibration with qualified standards for comparison. However, DNA or RNA quantities should be metrologically traceable to a reference (Bhat & Emslie, 2016; Whale et al., 2018). As of this writing, multiple nucleic acid quantitation methods have been developed, such as chemical analysis methods based on isotope-dilution mass spectrometry (IDMS), capillary electrophoresis (CE), and enumeration-based flow cytometric method (FCM) counting (Kwon, Jeong, Bae, Choi, & Yang, 2019; Yoo et al., 2016). These methods can be accurately calibrated with solutions of nucleic acids. In addition, an international comparison study was performed between national metrology institutes (NMI) using the dPCR method (Corbisier et al., 2012). Recently, the droplet digital PCR (ddPCR) method has emerged as a powerful analytical technique for clinical utility (Miotke, Lau, Rumma, & Ji, 2014; Pinheiro et al., 2012). For example, ddPCR can be used for the rapid enumeration of viral genomes and particles (Americo, Earl, & Moss, 2017; Hayden et al., 2013; Martinez-Hernandez et al., 2019).

In the present case of SARS-CoV-2, virus isolates or samples from infected patients have yet to be made available to the international public health community. In this study, we conclude that optimized conditions are required to increase the precision of ddPCR to develop reference materials with matrix conditions.

## Materials and Methods

### Construction of RNA standards

Full-length N and E genes of SARS-CoV-2 (GeneBank: MN908947) were synthesized and cloned in pET21a plasmid. The plasmids were amplified with the T7 promoter primer (5’-TAATACGACTCACTATAGGG-3’) and T7 terminator primer (5’-GCTAGTTATTGCTCAGCGG-3’). The amplicons were spectrophotometrically quantified at 260 nm. A total of 200 ng of the PCR product was used for in vitro transcription (MEGAscript T7 Transcription kit; Thermo Fisher Scientific, Waltham, MA, USA), which was performed at 37°C overnight in a 20 ul reaction mixture containing 2 ul of reaction buffer, 2 ul of each nucleoside triphosphate, and 2 ul of enzyme mix. The template DNAs were removed by digestion with 2 U of Turbo DNase I for 15 min at 37°C. The RNAs were precipitated by adding 6 ul 3 M sodium acetate and 150 ul of 98% ethanol which was followed by a subsequent incubation at −20°C for 30 min. After 15 min of centrifugation at 17,000 rpm, the supernatant was removed and 200 ul of 70% ethanol was added. After another 10-min centrifugation at 17,000 rpm, the supernatant was removed and the pellet was dissolved in 20 ul RNase-free H_2_O (Takara Bio Inc., Otsu, Shiga, Japan). Quantitation of the RNAs was performed spectrophotometrically at 260 nm. The measurement of the RNA concentrations was performed in duplicate, and the concentration was then converted to the molecule number (Fronhoffs et al., 2002).

### Cell and virus RNA extraction

The genomic RNA of clinical isolate SARS-CoV-2 (NCCP 43326) was obtained from the National Culture Collection for Pathogens (NCCP). The obtained RNA genomes were maintained at −80 °C and were used as a template for the assays. Vero E6 cells (ATCC CRL-1586) were purchased from the American Type Culture Collection (ATCC) and were maintained in Dulbecco’s modified Eagle medium (DMEM) supplemented with 2% (v/v) fetal bovine serum (FBS) and 1% antibiotic-antimycotic solution (Thermo Fisher Scientific, Waltham, MA, USA) at 37°C in a 5% humidified CO2 incubator. The SARS coronavirus HKU-39849 (GenBank: AY278491.2, provided by Dr. Malik Peiris of the University of Hong Kong) was inoculated into Vero E6 cells. Cytopathic effects were observed two days after inoculation and the viral titer was determined via a plaque assay. The viral RNA was extracted from the culture medium using the MagMAX-96 viral RNA isolation kit (Thermo Fisher Scientific, Waltham, MA, USA) according to the manufacturer’s instructions. Human hepatoma Huh-7 were obtained from the Japan Cell Research Bank (National Institutes of Biomedical Innovation, Health and Nutrition, Japan). The cells were maintained in Dulbecco’s modified Eagle’s medium (DMEM, HyClone, Logan, UT, USA) supplemented with 10% fetal bovine serum (FBS, HyClone, Logan, UT, USA) at 37 °C in a 5% CO_2_ incubator. The patient-derived isolate MERS-CoV strain KNIH/002_05_2015 (Kim et al., 2016) was obtained from KNIH and inoculated into cultured Huh-7 Cells. After 3 days of incubation, the viral RNA was extracted from the culture medium using the QIAamp viral RNA extraction Kit (Qiagen, Hilden, Germany) according to the manufacturer’s instructions. Human lung fibroblast MRC-5 cells were obtained from the American Type Culture Collection (ATCC). The culture medium for the cell line was Gibco Minimum Essential Media (MEM) supplemented with 10% heat-inactivated Fetal Bovine Serum (FBS), 1% Sodium Pyruvate, 1% penicillin, and 1% non-essential amino acids. The cell lines were cultured in a T75 flask in a humidified atmosphere of 5% CO_2_ at 37◻°C. The subcultures were produced with a ratio of 1:2 when the cell confluency reached 80%–90% every 3 or 4 days. HCoV OC43 (ATCC VR1588) was obtained from ATCC and was inoculated into a virus growth medium (MEM with 2% heat-inactivated FBS, 1% penicillin, and 1% non-essential amino acids) and propagated to the MRC-5 cells with 90% confluence. The flasks were maintained at 33◻°C in a humidified atmosphere of 5% CO_2_. When the cells reached 50%-60% confluence, the supernatant was harvested, filtered, and stored at −80◻°C until RNA extraction. Viral RNA was extracted using the viral RNA Minikit (QIAgen, Hilden, Germany) according to the manufacturer’s protocol.

### Clinical samples and RNA preparation

The clinical samples used in this study were collected from subjects according to registered protocols approved by the Institutional Review Board (IRB) of Jeonbuk National University Hospital, with all patients having signed written informed consent forms (IRB registration number: CUH 2020-02-050-008). The clinical characteristics of the patients are shown in Table 2. Upper respiratory tract specimens (naso- and oropharyngeal swabs) from COVID-19 patients were suspended in a transport medium (eNAT; COPAN, Murrieta, CA, USA) and stored at −80°C until use. The RNA extraction from the clinical samples was performed using the viral RNA Minikit (QIAgen, Hilden, Germany) according to the manufacturer’s protocol. The extracted viral RNA was used as a template for cDNA synthesis.

### Primer and probes

The primer-probe sets used in this study are listed in Table 1. For the detection of SARS-CoV-2, primer-probe sets of the Centers for Disease Control and Prevention (CDC, USA), The University of Hong Kong (HKU, Hong Kong), the National Institute of Infectious Disease Department of Virology III (NIID, Japan), and the National Institute of Health (NIH, Thailand) were selected according to laboratory guidance from WHO (Jung et al., 2020). The specific primer-probe sets for MERS-CoV and OC43 were used for the quantification of the viral RNA genomes (Corman et al., 2012; Vijgen et al., 2005). All primers and probes were synthesized by Neoprobe (Daejeon, Korea). All probes were labeled with the reporter molecule 6-carboxyfluorescein (FAM) at the 5’-end and with the quencher Black Hole Quencher 1 (BHQ-1) at the 3’-end.

**Table 1.**
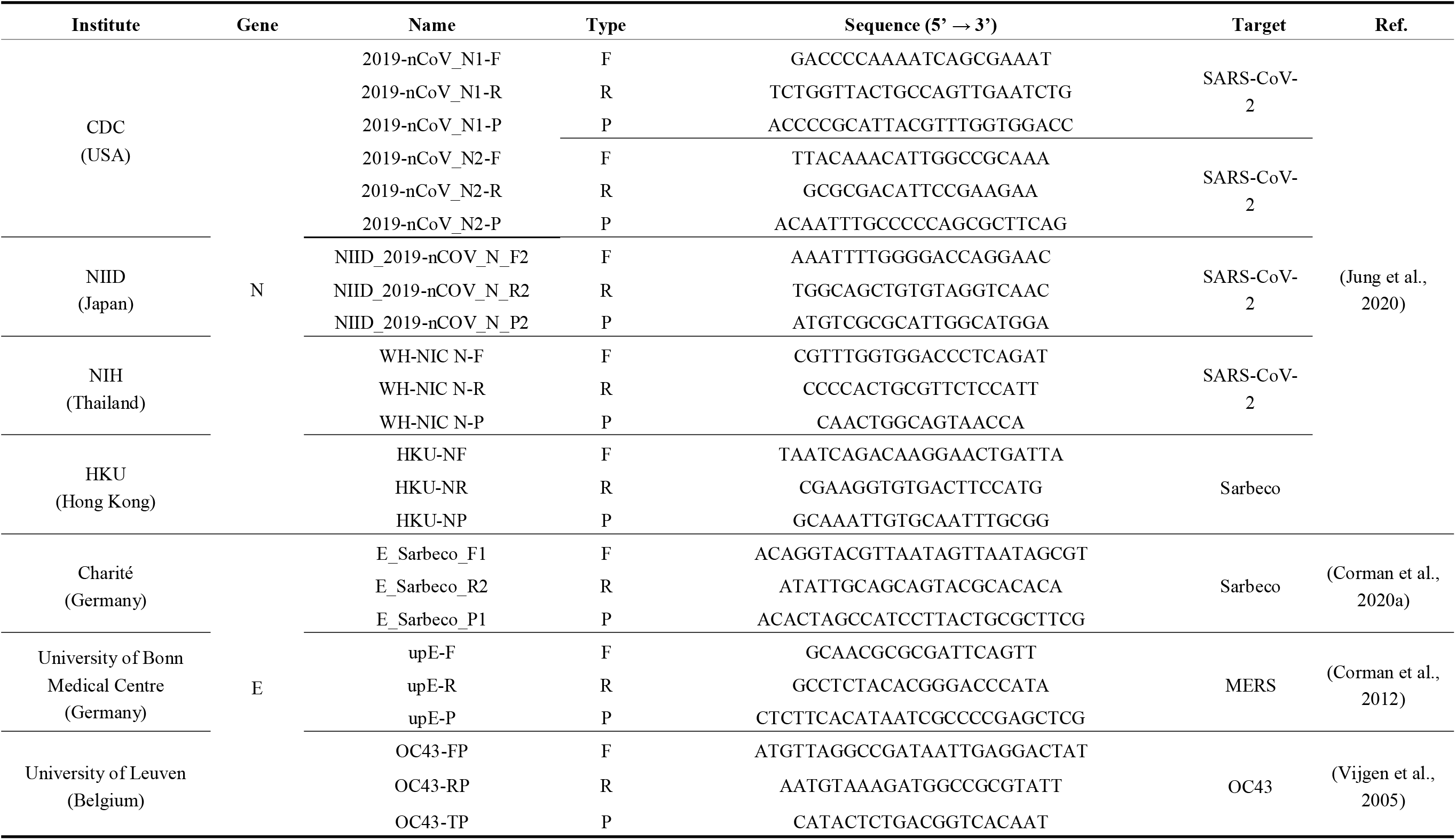
Information regarding the primers and probes used in this study.

### cDNA synthesis of viral RNA genomes

The cDNAs of the clinical and cultured (SARS-CoV-2, SARS-CoV, MERS-CoV, and OC43) viral RNA genomes were synthesized using the LunaScript RT SuperMix Kit (New England Biolabs, Ipswich, MA, USA). The reaction mixture had a volume of 20◻μL and comprised of 4◻μL of 5X LunaScript RT supermix, 1◻μL of cDNA template, and 15 μL of distilled water. The reactions were carried out according to the manufacturer’s instructions. The synthesized cDNAs were serially diluted with distilled water and used as templates for qPCR and ddPCR without purification.

### qPCR and droplet digital PCR measurement

The templates were serially diluted to few copy numbers/μL. RT-qPCR analysis was performed on the StepOne and StepOnePlus Real-Time PCR system (Thermo Fisher Scientific, Waltham, MA, USA). The reaction mixture had a volume of 25◻μL and comprised of 12.5◻μL of maxima Probe/ROX qPCR master mix (2X) (Thermo Fisher Scientific, Waltham, MA, USA), 1◻μL of cDNA template, 1◻μL of 10◻μM forward primer, 1◻μL of 10◻μM reverse primer, 1◻μL of 5 μM probe labeled with FAM, and 8.5 μL of distilled water. The ddPCR analysis was performed using a QX200 system (BioRad Laboratories, Hercules, CA, USA). The reaction mixture had a volume of 20◻μL and comprised of 10◻μL of ddPCR supermix for probes (BioRad Laboratories, Hercules, CA, USA), 1◻μL of cDNA template, 1◻μL of 10◻μM forward primer, 1◻μL of 10◻μM reverse primer, 1◻μL of 5 μM probe labeled with FAM, and 6 μL of distilled water. All valid copy numbers were selected according to the manufacturer’s instructions (BioRad, 2015).

### Data analysis

The qPCR data were initially analyzed using the StepOne software (Thermo Fisher Scientific, Waltham, MA, USA). Raw data (i.e., the fluorescence values for C_T_) were exported from the StepOne software to Microsoft Excel 2016. A series of diluted templates were used to determine the C_T_ value, which can establish a standard curve for evaluating the reaction efficiency. Droplet fluorescence data were initially analyzed using the QuantaSoft software (BioRad Laboratories, Hercules, CA, USA). Raw data (i.e., the fluorescence values for the droplets) were exported from the QuantaSoft software to Microsoft Excel 2016. Depending on the separation of positive and negative droplets, an objective separation value k was automatically calculated.

## Results

### Quantification of viral and IVT RNAs

In this study, the primer-probe sets of each E gene were used to quantify the viral genomic RNA of SARS-CoV-2 and related coronaviruses (Table 1) (Corman et al., 2012, 2020a; Jung et al., 2020; Vijgen et al., 2005). The clinical characteristics of confirmed COVID-19 patients with symptoms are summarized in Table 2.

**Table 2.** Clinical characteristics of laboratory-confirmed patients with COVID-19.

|  | Patient 1 | Patient 2 | Patient 3 | Patient 4 | Patient 5 |
| --- | --- | --- | --- | --- | --- |
| Age (years)/Sex | 72/M | 57/M | 72/F | 48/M | 51/F |
| Fever | - | + | + | + | + |
| Chills | - | + | - | - | - |
| Cough | + | + | + | - | + |
| Dyspnea | - | + | + | - | + |
| Diarrhea | - | - | - | + | - |
| Laboratory findings |  |  |  |  |  |
| WBC | 3,570 | 3,710 | 6,720 | 4,730 | 5,820 |
| Hemoglobin | 10.3 | 12.9 | 16.1 | 12.9 | 10.8 |
| Platelet | 157,000 | 42,000 | 190,000 | 316,000 | 175,000 |
| AST | 23 | 57 | 32 | 34 | 43 |
| ALT | 22 | 55 | 13 | 38 | 18 |
| Creatinine | 0.83 | 0.77 | 0.55 | 0.31 | 0.38 |
| Used drug |  |  |  |  |  |
| Lopinavir/ritonavir | + | + | + | + | + |
| Hydroxychloroquine | + | + | + | + | + |
| Outcome | Cured | Cured | Cured | Cured | Cured |

The copy numbers of SARS-CoV-2, SARS-CoV, MERS, and OC43 viral RNA were measured using ddPCR, and the measured values of viral genomic RNA were 8.04 × 10^5^, 6.52 × 10^5^, 9.37 × 10^4^ and 9.57 × 10^4^ copies/μL, respectively (Table 3). To assess the quantification of viral genomic RNA and partial synthetic viral RNA, *in vitro* transcript (IVT) N and E gene RNA were measured via qPCR and ddPCR. The copy numbers of N IVT RNA and E IVT RNA were measured as 1.8 × 10^9^ and 3.47 × 10^9^, respectively. The amplification efficiencies were calculated from standard curves of SARS-CoV-2 viral RNA and IVT RNAs.

**Table 3.** R^2^ values calculated with mean C_T_ and copy number quantified via qPCR and ddPCR.

| Target<br>Dilution factor | E_Sarbeco |  |  |  |
| --- | --- | --- | --- | --- |
|  | SARS-CoV-2 |  | SARS-CoV |  |
| | qPCR ($C_T$) | ddPCR (copy/μl) | qPCR ($C_T$) | ddPCR (copy/μl) |
| $10^{-1}$ | 19.81 | 98533 | 19.96 | 90000 |
| $10^{-2}$ | 23.09 | 9780 | 23.34 | 8747 |
| $10^{-3}$ | 26.79 | 804 | 27.30 | 652 |
| $10^{-4}$ | 31.20 | 46 | 30.77 | 57 |
| Efficiency (%) | 83.65 |  | 88.30 |  |
| $R^2$ | 0.9956 | | 0.9991 | |
| Target<br>Dilution factor | E_upE |  | E_OC43 |  |
|  | MERS |  | OC43 |  |
| | qPCR ($C_T$) | ddPCR (copy/μl) | qPCR ($C_T$) | ddPCR (copy/μl) |
| 1 | 24.46 | 86400 | 24.69 | 99800 |
| $10^{-1}$ | 27.73 | 8980 | 27.93 | 10887 |
| $10^{-2}$ | 31.06 | 937 | 31.46 | 957 |
| $10^{-3}$ | 35.11 | 74 | 35.16 | 93 |
| Efficiency (%) | 92.01 |  | 93.24 |  |
| $R^2$ | 0.9972 | | 0.9991 | |
| Target | 2019-nCoV_N2 |  | E_Sarbeco |  |

| Dilution factor | IVT N gene |  | IVT E gene |  |
| --- | --- | --- | --- | --- |
|  | qPCR (C<sub>T</sub>) | ddPCR (copy/μl) | qPCR (C<sub>T</sub>) | ddPCR (copy/μl) |
| 10<sup>-4</sup> | 19.61 | 6794000 | 20.94 | 20000000 |
| 10<sup>-5</sup> | 24.03 | 22280 | 24.97 | 43490 |
| 10<sup>-6</sup> | 28.09 | 1803 | 28.90 | 3470 |
| 10<sup>-7</sup> | 31.85 | 154 | 32.49 | 318 |
| Efficiency (%) | 75.88 |  | 81.71 |  |
| R<sup>2</sup> | 0.9987 |  | 0.9993 |  |

\* The initial concentrations of SARS-CoV-2, SARS-CoV, MERS, OC43 viral RNA, and IVT N gene and E gene RNAs were measured using a Quantus fluorometer and the measured values of each RNA were 78.3, 35.7, 30.8, 66.2, 91.4, and 50.9 ng/μL, respectively.

### IVT RNAs as a standard for RT-qPCR

The R^2^ values of viral RNA and IVT N IVT RNA were calculated as 0.9973 and 0.9987, respectively, whereas the R^2^ values of viral RNA and IVT E IVT RNA were calculated as 0.9956 and 0.9993, respectively (Tables 3 and 4). These values indicate that the standard curves of both viral and IVT RNAs are linear and accurate. The amplification efficiency of both viral and IVT RNAs were calculated from the slopes of the standard curves. The efficiencies of both viral and IVT RNAs according to the target genes were similar (Tables 3 and 4). The similar amplification efficiencies show that the quantified IVT RNA can be used as a standard for RT-qPCR measurement of viral RNA.

**Table 4.** Comparative analysis of C_T_ values and copy numbers obtained via qPCR and ddPCR.

| Dilution<br>factor | 2019-nCoV_N1 |  | 2019-nCoV_N2 |  | NIID_2019-nCoV_N |  | WH-NIC_N |  | HKU_N |  |
| --- | --- | --- | --- | --- | --- | --- | --- | --- | --- | --- |
|  | qPCR | ddPCR | qPCR | ddPCR | qPCR | ddPCR | qPCR | ddPCR | qPCR | ddPCR |
| 10<sup>-2</sup> | 18.89 | 6806000 | 18.87 | 175333 | 18.14 | 163000 | 21.40 | 6797333 | 20.79 | 165400 |
| 10<sup>-3</sup> | 22.16 | 21173 | 22.19 | 20340 | 21.63 | 15173 | 24.65 | 19993 | 24.31 | 18473 |
| 10<sup>-4</sup> | 25.99 | 1907 | 26.06 | 1659 | 25.40 | 1177 | 28.52 | 1599 | 28.09 | 1553 |
| 10<sup>-5</sup> | 30.66 | 135 | 30.60 | 132 | 29.93 | 60 | 33.11 | 105 | 32.68 | 105 |
| 10<sup>-6</sup> | 34.12 | 11 | 34.57 | 8 | 34.91 | 7 | 36.71 | 5 | 36.71 | 12 |
| Efficiency (%) | 80.58 |  | 78.28 |  | 73.40 |  | 80.26 |  | 77.28 |  |
| R<sup>2</sup> | 0.9968 |  | 0.9973 |  | 0.9943 |  | 0.9971 |  | 0.9978 |  |

\* A copy number quantified as a target of Germany E gene using ddPCR

### Comparison of qPCR and ddPCR assays for COVID-19 clinical samples

According to the qPCR and ddPCR results, all patient samples were positive when we used 0.1 ng of extracted RNA for the template. The non-template control did not produce any signals from both the qPCR and ddPCR assays. The result of the serially diluted sample showed that the C_T_ value was undetermined. On the other hand, all ddPCR signals were detected in all of the diluted samples (Table 5). Furthermore, the positive copies (P2 for 0.67 and P3 for 0.60) from the two samples were exceptionally low compared to the other specimens, but still only produced positive signals in the ddPCR results. This suggests that the ddPCR method can be applied to detect ultra-low amounts of sample with high sensitivity.

**Table 5.** C_T_ values and copy numbers of patient samples via qPCR and ddPCR.

| Dilution<br>factor | Template | 2019-nCoV_N2 primer-probe set |  |  |  |  |  |  |  |  |  |
| --- | --- | --- | --- | --- | --- | --- | --- | --- | --- | --- | --- |
|  |  | P1<sup>*</sup> |  | P2<sup>*</sup> |  | P3<sup>*</sup> |  | P4<sup>*</sup> |  | P5<sup>*</sup> |  |
|  |  | qPCR<br>(Ct) | ddPCR<br>(copy/μl) | qPCR<br>(Ct) | ddPCR<br>(copy/μl) | qPCR<br>(Ct) | ddPCR<br>(copy/μl) | qPCR<br>(Ct) | ddPCR<br>(copy/μl) | qPCR<br>(Ct) | ddPCR<br>(copy/μl) |
| 10<sup>-1</sup> |  | 26.87 | 1051 | 35.03 | 7.20 | 35.15 | 2.40 | Undetermined | 0 | 35.05 | 7.53 |
| 10<sup>-2</sup> |  | 30.52 | 82 | 37.03 | 1.33 | Undetermined | 0.73 | Undetermined | 0.53<sup>**</sup> | Undetermined | 1.20 |
| 10<sup>-3</sup> |  | 33.81 | 5.93 | Undetermined | 0.67 | Undetermined | 0.60 | Undetermined | 0 | Undetermined | 1.07 |

<sup>\*</sup> The initial concentrations of the viral RNA templates were 46.9, 21.1, 27.5, 17.2, and 15.9 ng/μL, respectively.
<sup>\*\*</sup> Only one of the triplicate was positive (single droplet).

## Discussion

The outbreak of COVID-19 is a pandemic threat caused by the emergence of SARS-CoV-2, a newly discovered human coronavirus (Wu et al., 2020; Zhou et al., 2020). SARS-CoV-2 has four structural proteins, known as the S (spike), E (envelope), M (membrane), and N (nucleocapsid) proteins (Walls et al., 2020). The SARS-CoV-2 S protein binds with angiotensin-converting enzyme 2 (ACE2) with high-affinity, which leads to clinical features of pneumonia in many patients (Wrapp et al., 2020; Zhu et al., 2020). SARS-CoV-2 is highly contagious and has quickly spread across the world, which further emphasizes the essential role of diagnostics in the control of communicable diseases (Winter & Hegde, 2020).

Currently, a reliable test (also the most widely used) for detecting acute infections of SARS-CoV-2 is the reverse transcription-quantitative polymerase chain reaction (RT-qPCR) method (Lu et al., 2020). Multiple national laboratories have developed assays targeting conserved regions that could potentially detect pan-sarbecovirus (Chu et al., 2020; Corman et al., 2020b). A previous study provided information such as the specific primer and probe sequence, thermal profile, and reagents for RT-qPCR to optimize the reaction conditions for the detection of SARS-CoV-2 RNA (Bustin, 2020). ddPCR is a method that improves upon conventional PCR methods that can individually amplify and directly quantify pathogen RNA or DNA (Li et al., 2018). Several studies have compared RT-qPCR and ddPCR for the quantification of viral genomes (Abachin et al., 2018; Dobnik et al., 2019; Kiselinova et al., 2014).

In the present study, SARS-CoV-2 RNA concentrations were measured using the qPCR and ddPCR methods (Table 3). The qPCR results showed that the C_T_ value of a given template was greatly dependent on the individual primer-probe sets. However, the copy numbers determined via ddPCR were stable and reliable in the quantification of low-viral RNA samples, regardless of the primer-probe sets. A previous study reported that qPCR and ddPCR results were consistent for 95 positive samples and that there was a high correlation between the C_T_ values of qPCR and the copy number values of ddPCR among patient samples (Yu et al., 2020). Although the C_T_ values of each primer-probe set exhibited variance, the copy numbers from ddPCR showed that N genes are more abundant than E genes. These findings are also consistent with another previous study, indicating N genes are good targets for diagnostics (Kim et al., 2020).

The qPCR method requires the use of a standard curve to obtain quantitative measurements of an unknown sample (Adamski, Gumann, & Baird, 2014). However, in the case of ddPCR, the distribution of target-specific amplification into partitions is calculated using the Poisson distribution, enabling the absolute quantification of the target gene based on the ratio of positive against all partitions at the end of the reaction without dependence on an external standard (Gutiérrez-Aguirre, Rački, Dreo, & Ravnikar, 2015). In addition to these advantages, our results revealed that ddPCR is a more sensitive and accurate method for SARS-CoV-2 detection in low viral copies (Table 5). Further experiments should be performed even if patients are cured of COVID-19, and such tests should be conducted using ddPCR to lower the limit of detection (LOD) to be able to detect low amounts of viral RNA.

The Coronavirus Standards Working Group led by the Joint Initiative for Metrology in Biology (JIMB) is conducting international collaborative experiments to develop guidelines for the purpose of evaluating and establishing the availability of common and appropriate standards, diverse reference materials, validation tests, and reference measurement protocols that can enable reliable COVID-19 testing (“Coronavirus Standards Working Group — The Joint Initiative for Metrology in Biology,” in press; “ISO-COVID-19 response: freely available ISO standards,” in press). Our results serve as a guide for the standardization of analytical methods and the development of reference materials that are required for accurate COVID-19 diagnosis based on both RT-qPCR and ddPCR. Consequently, ddPCR-based diagnostics can serve as a companion method to the current standard RT-qPCR to provide sensitive and accurate quantitative results for emergency testing that can be adopted for clinical use.

## Conclusion

Our study demonstrated that CT values of samples from qPCR assays were significantly variable depending on the sequence of primer and probe sets, whereas copy numbers of samples from ddPCR assays were hardly affected by the sequence of primer and probe sets. These results indicate that ddPCR can be performed with sub-optimal primer-probe sets without the loss of sensitivity. A comparison of the standard curves of viral RNA and IVT RNA showed that IVT RNA quantified via ddPCR could be used as a standard for the absolute quantification of qPCR assays. The results from the clinical samples showed that the ddPCR method could result in highly sensitive and quantitative diagnostics. Although qPCR produced different CT values using different primer-probes sets, the ddPCR results obtained using different primer-probe sets were compatible with each other. Therefore, the deployment of ddPCR-based diagnostics could produce highly sensitive and quantitative results that are compatible between different laboratories.

## Author Contributions

Conceptualization, S. K. and H. M. Y.; methodology, S. K. and H. M. Y.; validation, S. K. and H. M. Y.; formal analysis, C. P.; investigation, C. P., J. L., Z. U. H., K. B. K., S. J. K., H. G. K., E. C. P., G.-S. P., D. P., S.-H. B., D. K., D. P., J. L., S. J., S. K., and C.-S. L.; resources, C. P., J. L., Z. U. H., K. B. K., E. C. P., S.-H. B., D. P., J. L., S. J., S. K., and C.-S. L.; data curation, S. K. and H. M. Y.; writing—original draft preparation, C. P., Z. U. H., K. B. K., E. C. P., S. K., C.-S. L., S. K. and H. M. Y.; writing—review and editing, S. K. and H. M. Y.; visualization, C. P., S. K. and H. M. Y.; supervision, S. K. and H. M. Y.; project administration, S. K..; funding acquisition, S. K.

## Acknowledgements

This work was supported by National Research Council of Science and Technology grant (Grant No. CRC◻16◻01◻KRICT). This research was also supported by the “Development of recombinant coronavirus reference materials”, grant number KRISS-2020-GP2020-0003-08, and “Development of Measurement Standards and Technology for Biomaterials and Medical Convergence”, grant number KRISS-2020-GP2020-0004 programs, funded by the Korea Research Institute of Standards and Science. ORCID ID (Hee Min Yoo: 0000-0002-5951-2137; Seil Kim: 0000-0003-3465-7118; Changwoo Park: 0000-0001-7083-9037; Jina Lee: 0000-0002-3661-3701; Zohaib Ul Hassan: 0000-0002-4982-2262; Dongju Park: 0000-0001-6516-6755

## References

Abachin, E., Convers, S., Falque, S., Esson, R., Mallet, L., & Nougarede, N. 2018. Comparison of reverse-transcriptase qPCR and droplet digital PCR for the quantification of dengue virus nucleic acid. Biologicals, 52(February): 49–54.

Adamski, M. G., Gumann, P., & Baird, A. E. 2014. A method for quantitative analysis of standard and high-throughput qPCR expression data based on input sample quantity. PLoS ONE, 9(8).

Americo, J. L., Earl, P. L., & Moss, B. 2017. Droplet digital PCR for rapid enumeration of viral genomes and particles from cells and animals infected with orthopoxviruses. Virology, 511(August): 19–22.

Ashour, H. M., Elkhatib, W. F., Rahman, M. M., & Elshabrawy, H. A. 2020. Insights into the recent 2019 novel coronavirus (Sars-coV-2) in light of past human coronavirus outbreaks. Pathogens, 9(3): 1–15.

Bhat, S., & Emslie, K. R. 2016. Digital polymerase chain reaction for characterisation of DNA reference materials. Biomolecular Detection and Quantification, 10: 47–49.

BioRad. 2015. Rare Mutation Detection Best Practices Guidelines.

Bustin, S. A. 2020. RT-qPCR Testing of SARS-CoV-2◻ A Primer.

Chu, D. K. W., Pan, Y., Cheng, S. M. S., Hui, K. P. Y., Krishnan, P., Liu, Y., Ng, D. Y. M., Wan, C. K. C., Yang, P., Wang, Q., Peiris, M., & Poon, L. L. M. 2020. Molecular Diagnosis of a Novel Coronavirus (2019-nCoV) Causing an Outbreak of Pneumonia. Clinical Chemistry, 66(4): 549–555.

Corbisier, P., Vincent, S., Schimmel, H., Kortekaas, A.-M., Trapmann, S., Burns, M., Bushell, C., Akgoz, M., Akyürek, S., Dong, L., Fu, B., Zhang, L., Wang, J., Pérez Urquiza, M., Bautista, J. L., Garibay, A., Fuller, B., Baoutina, A., Partis, L., Emslie, K., et al. 2012. CQM-K86/P113.1: Relative quantification of genomic DNA fragments extracted from a biological tissue. Metrologia, 49(1A): 08002–08002.

Corman, V. M., Eckerle, I., Bleicker, T., Zaki, A., Landt, O., Eschbach-Bludau, M., van Boheemen, S., Gopal, R., Ballhause, M., Bestebroer, T. M., Muth, D., Müller, M. A., Drexler, J. F., Zambon, M., Osterhaus, A. D., Fouchier, R. M., & Drosten, C. 2012. Detection of a novel human coronavirus by real-time reverse-transcription polymerase chain reaction. Eurosurveillance, 17(39).

Corman, V. M., Landt, O., Kaiser, M., Molenkamp, R., Meijer, A., Chu, D. K. W., Bleicker, T., Brünink, S., Schneider, J., Schmidt, M. L., Mulders, D. G. J. C., Haagmans, B. L., Van Der Veer, B., Van Den Brink, S., Wijsman, L., Goderski, G., Romette, J. L., Ellis, J., Zambon, M., Peiris, M., et al. 2020a. Detection of 2019 novel coronavirus (2019-nCoV) by real-time RT-PCR. Eurosurveillance, 25(3).

Corman, V. M., Landt, O., Kaiser, M., Molenkamp, R., Meijer, A., Chu, D. K. W., Bleicker, T., Brünink, S., Schneider, J., Schmidt, M. L., Mulders, D. G. J. C., Haagmans, B. L., Van Der Veer, B., Van Den Brink, S., Wijsman, L., Goderski, G., Romette, J. L., Ellis, J., Zambon, M., Peiris, M., et al. 2020b. Detection of 2019 novel coronavirus (2019-nCoV) by real-time RT-PCR. Eurosurveillance, 25(3): 1–8.

Coronaviridae Study Group of the International Committee on Taxonomy of Viruses. 2020. The species Severe acute respiratory syndrome-related coronavirus: classifying 2019-nCoV and naming it SARS-CoV-2. Nature microbiology, 5(March).

Coronavirus Standards Working Group — The Joint Initiative for Metrology in Biology. in press. Retrieved April 27, 2020, from https://jimb.stanford.edu/covid-19-standards

da Costa, V. G., Moreli, M. L., & Saivish, M. V. 2020. The emergence of SARS, MERS and novel SARS-2 coronaviruses in the 21st century. Archives of virology, (0123456789).

Dobnik, D., Kogovšek, P., Jakomin, T., Košir, N., Žnidarič, M. T., Leskovec, M., Kaminsky, S. M., Mostrom, J., Lee, H., & Ravnikar, M. 2019. Accurate quantification and characterization of adeno-associated viral vectors. Frontiers in Microbiology, 10(JULY).

Fronhoffs, S., Totzke, G., Stier, S., Wernert, N., Rothe, M., Brüning, T., Koch, B., Sachinidis, A., Vetter, H., & Ko, Y. 2002. A method for the rapid construction of cRNA standard curves in quantitative real-time reverse transcription polymerase chain reaction. Molecular and Cellular Probes, 16(2): 99–110.

Fung, T. S., & Liu, D. X. 2019. Human Coronavirus: Host-Pathogen Interaction. Annual Review of Microbiology, 73(1): 529–557.

Ge, X. Y., Li, J. L., Yang, X. Lou, Chmura, A. A., Zhu, G., Epstein, J. H., Mazet, J. K., Hu, B., Zhang, W., Peng, C., Zhang, Y. J., Luo, C. M., Tan, B., Wang, N., Zhu, Y., Crameri, G., Zhang, S. Y., Wang, L. F., Daszak, P., & Shi, Z. L. 2013. Isolation and characterization of a bat SARS-like coronavirus that uses the ACE2 receptor. Nature, 503(7477): 535–538.

Gorbalenya, A. E. 2020. Severe acute respiratory syndrome-related coronavirus – The species and its viruses, a statement of the Coronavirus Study Group. BioRxiv, 2020.02.07.937862.

Gutiérrez-Aguirre, I., Rački, N., Dreo, T., & Ravnikar, M. 2015. Droplet digital PCR for absolute quantification of pathogens. Methods in Molecular Biology, 1302: 331–347.

Hayden, R. T., Gu, Z., Ingersoll, J., Abdul-Ali, D., Shi, L., Pounds, S., & Caliendo, A. M. 2013. Comparison of droplet digital PCR to real-time PCR for quantitative detection of cytomegalovirus. Journal of Clinical Microbiology, 51(2): 540–546.

Hindson, B. J., Ness, K. D., Masquelier, D. A., Belgrader, P., Heredia, N. J., Makarewicz, A. J., Bright, I. J., Lucero, M. Y., Hiddessen, A. L., Legler, T. C., Kitano, T. K., Hodel, M. R., Petersen, J. F., Wyatt, P. W., Steenblock, E. R., Shah, P. H., Bousse, L. J., Troup, C. B., Mellen, J. C., Wittmann, D. K., et al. 2011. High-Throughput Droplet Digital PCR System for Absolute Quantitation of DNA Copy Number. Analytical Chemistry, 83(22): 8604–8610.

Hindson, C. M., Chevillet, J. R., Briggs, H. A., Gallichotte, E. N., Ruf, I. K., Hindson, B. J., Vessella, R. L., & Tewari, M. 2013. Absolute quantification by droplet digital PCR versus analog real-time PCR. Nature Methods, 10(10): 1003–1005.

ISO-COVID-19 response: freely available ISO standards. in press. Retrieved April 27, 2020, from https://www.iso.org/covid19

Jung, Y. J., Park, G.-S., Moon, J. H., Ku, K., Beak, S.-H., Kim, S., Park, E. C., Park, D., Lee, J.-H., Byeon, C. W., Lee, J. J., Maeng, J., Kim, S. J., Kim, S. Il, Kim, B.-T., Lee, M. J., & Kim, H. G. 2020. Comparative analysis of primer-probe sets for the laboratory confirmation of SARS-CoV-2. BioRxiv, 2020.02.25.964775.

Kim, D., Lee, J.-Y., Yang, J.-S., Kim, J. W., Kim, V. N., & Chang, H. 2020. The Architecture of SARS-CoV-2 Transcriptome. Cell, 181(0): 1–8.

Kim, D. W., Kim, Y. J., Park, S. H., Yun, M. R., Yang, J. S., Kang, H. J., Han, Y. W., Lee, H. S., Kim, H. M., Kim, H., Kim, A. R., Heo, D. R., Kim, S. J., Jeon, J. H., Park, D., Kim, J. A., Cheong, H. M., Nam, J. G., Kim, K., & Kim, S. S. 2016. Variations in spike glycoprotein gene of MERS-coV, South Korea, 2015. Emerging Infectious Diseases, 22(1): 100–104.

Kiselinova, M., Pasternak, A. O., De Spiegelaere, W., Vogelaers, D., Berkhout, B., & Vandekerckhove, L. 2014. Comparison of droplet digital PCR and seminested real-time PCR for quantification of cell-associated HIV-1 RNA. PLoS ONE, 9(1): 1–8.

Kwon, H. J., Jeong, J. S., Bae, Y. K., Choi, K., & Yang, I. 2019. Stable isotope labeled DNA: A new strategy for the quantification of total dna using liquid chromatography-mass spectrometry. Analytical Chemistry.

Li, H., Bai, R., Zhao, Z., Tao, L., Ma, M., Ji, Z., Jian, M., Ding, Z., Dai, X., Bao, F., & Liu, A. 2018. Application of droplet digital PCR to detect the pathogens of infectious diseases. Bioscience Reports, 38(6): 1–8.

Li, W., Wong, S.-K., Li, F., Kuhn, J. H., Huang, I.-C., Choe, H., & Farzan, M. 2006. Animal Origins of the Severe Acute Respiratory Syndrome Coronavirus: Insight from ACE2-S-Protein Interactions. Journal of Virology, 80(9): 4211–4219.

Lu, R., Wu, X., Wan, Z., Li, Y., Jin, X., & Zhang, C. 2020. A Novel Reverse Transcription Loop-Mediated Isothermal Amplification Method for Rapid Detection of SARS-CoV-2. International Journal of Molecular Sciences, 21(8): 2826.

Martinez-Hernandez, F., Garcia-Heredia, I., Gomez, M. L., Maestre-Carballa, L., Martínez, J. M., & Martinez-Garcia, M. 2019. Droplet digital PCR for estimating absolute abundances of widespread pelagibacter viruses. Frontiers in Microbiology, 10(JUN): 1–13.

Miotke, L., Lau, B. T., Rumma, R. T., & Ji, H. P. 2014. High sensitivity detection and quantitation of DNA copy number and single nucleotide variants with single color droplet digital PCR. Analytical Chemistry, 86(5): 2618–2624.

Pinheiro, L. B., Coleman, V. A., Hindson, C. M., Herrmann, J., Hindson, B. J., Bhat, S., & Emslie, K. R. 2012. Evaluation of a droplet digital polymerase chain reaction format for DNA copy number quantification. Analytical Chemistry, 84(2): 1003–1011.

Vijgen, L., Keyaerts, E., Moës, E., Maes, P., Duson, G., & Van Ranst, M. 2005. Development of one-step, real-time, quantitative reverse transcriptase PCR assays for absolute quantitation of human coronaviruses OC43 and 229E. Journal of Clinical Microbiology, 43(11): 5452–5456.

Walls, A. C., Park, Y. J., Tortorici, M. A., Wall, A., McGuire, A. T., & Veesler, D. 2020. Structure, Function, and Antigenicity of the SARS-CoV-2 Spike Glycoprotein. Cell, 1–12.

Whale, A. S., Jones, G. M., Pavšič, J., Dreo, T., Redshaw, N., Akyurek, S., Akgoz, M., Divieto, C., PaolaSassi, M., He, H.-J., Cole, K. D., Bae, Y.-K., Park, S.-R., Deprez, L., Corbisier, P., Garrigou, S., Taly, V., Larios, R., Cowen, S., O’Sullivan, D. M., et al. 2018. Assessment of Digital PCR as a Primary Reference Measurement Procedure to Support Advances in Precision Medicine. Clinical Chemistry, 64(9): 1296–1307.

WHO | Middle East respiratory syndrome coronavirus (MERS-CoV). in press. Retrieved March 23, 2020, from https://www.who.int/emergencies/mers-cov/en/

WHO | Summary of probable SARS cases with onset of illness from 1 November 2002 to 31 July 2003. 2015. WHO.

Winter, A. K., & Hegde, S. T. 2020. The important role of serology for COVID-19 control. The Lancet Infectious Diseases, 0(0).

World Health Organization (WHO). 2020. Novel Coronavirus (2019-nCoV) Situation Report −1 21 January 2020. WHO Bulletin, (JANUARY): 1–7.

Wrapp, D., Wang, N., Corbett, K. S., Goldsmith, J. A., Hsieh, C. L., Abiona, O., Graham, B. S., & McLellan, J. S. 2020. Cryo-EM structure of the 2019-nCoV spike in the prefusion conformation. Science, 367(6483): 1260–1263.

Wu, F., Zhao, S., Yu, B., Chen, Y. M., Wang, W., Song, Z. G., Hu, Y., Tao, Z. W., Tian, J. H., Pei, Y. Y., Yuan, M. L., Zhang, Y. L., Dai, F. H., Liu, Y., Wang, Q. M., Zheng, J. J., Xu, L., Holmes, E. C., & Zhang, Y. Z. 2020. A new coronavirus associated with human respiratory disease in China. Nature, 579(7798): 265–269.

Wu, K., Peng, G., Wilken, M., Geraghty, R. J., & Li, F. 2012. Mechanisms of host receptor adaptation by severe acute respiratory syndrome coronavirus. Journal of Biological Chemistry, 287(12): 8904–8911.

Yoo, H. B., Park, S. R., Dong, L., Wang, J., Sui, Z., Pavsic, J., Milavec, M., Akgoz, M., Mozioglu, E., Corbisier, P., Janka, M., Cosme, B., Cavalcante, J. J. D. V., Flatshart, R. B., Burke, D., Forbes-Smith, M., McLaughlin, J., Emslie, K., Whale, A. S., Huggett, J. F., et al. 2016. International comparison of enumeration-based quantification of DNA copy-concentration using flow cytometric counting and digital polymerase chain reaction. Analytical Chemistry, 88(24): 12169–12176.

Yu, F., Yan, L., Wang, N., Yang, S., Wang, L., Tang, Y., Gao, G., Wang, S., Ma, C., Xie, R., Wang, F., Tan, C., Zhu, L., Guo, Y., & Zhang, F. 2020. Quantitative Detection and Viral Load Analysis of SARS-CoV-2 in Infected Patients. Clinical Infectious Diseases.

Zhou, P., Yang, X. Lou, Wang, X. G., Hu, B., Zhang, L., Zhang, W., Si, H. R., Zhu, Y., Li, B., Huang, C. L., Chen, H. D., Chen, J., Luo, Y., Guo, H., Jiang, R. Di, Liu, M. Q., Chen, Y., Shen, X. R., Wang, X., Zheng, X. S., et al. 2020. A pneumonia outbreak associated with a new coronavirus of probable bat origin. Nature, 579(7798): 270–273.

Zhu, N., Zhang, D., Wang, W., Li, X., Yang, B., Song, J., Zhao, X., Huang, B., Shi, W., Lu, R., Niu, P., Zhan, F., Ma, X., Wang, D., Xu, W., Wu, G., Gao, G. F., & Tan, W. 2020. A novel coronavirus from patients with pneumonia in China, 2019. New England Journal of Medicine, 382(8): 727–733.

